# The Secreted Tyrosine Phosphatase PtpA promotes *Staphylococcus aureus* Intramacrophagic Survival Through Decrease of the SUMOylation Host Response

**DOI:** 10.1101/2022.08.31.506036

**Authors:** Nadhuma Youssouf, Marianne Martin, Markus Bischoff, Philippe Soubeyran, Laila Gannoun-Zaki, Virginie Molle

**Affiliations:** Laboratory of Pathogen Host Interactions, Université de Montpellier, CNRS, UMR 5235, Montpellier, France; (N.Y), (M.M), (L.G-Z) and (V.M); Institute of Medical Microbiology and Hygiene, University of Saarland, Homburg/Saar, Germany; (M.B); Centre de Recherche en Cancérologie de Marseille (CRCM), INSERM U1068, CNRS UMR 7258, Aix-Marseille, Université and Institut Paoli-Calmettes, Parc Scientifique et Technologique de Luminy, Marseille, France; (P.S)

**Keywords:** *Staphylococcus aureus*, SUMOylation, secreted phosphatase, PtpA, macrophage survival

## Abstract

*S. aureus* is a human pathogen that is extremely adaptable and is the cause of a variety of nosocomial and community-acquired infectious illnesses. During infection, *S. aureus* affect the host cell in many ways to enable its own multiplication, spread, and evasion of host immune defense. One of *S. aureus* mechanism to survive is to inhibit the SUMOylation of host proteins in order to increase its intracellular survival and persistence for longer period of time. Here, we show that the reduction in the levels of cellular SUMO-conjugated proteins is associated to the PtpA secreted virulence factor, which results in a reduction of Ubc9 protein level, the essential enzyme of the SUMOylation modification. In addition, we demonstrated that the critical residue D120A, essential for PtpA phosphatase activity, is required. This study shows for the first time that the secreted phosphatase PtpA impedes the host SUMOylation response, thus promoting *S. aureus* survival at long-term infection.

## 1. Introduction

Pathogenic bacteria affect the host cell in many ways during infection to enable their own multiplication, spread, and evasion of host immune defense [1]. Post-translational modifications (PTMs), which are essential for controlling the location, activity, and interaction of cellular components such lipids, cofactors, nucleic acids, and other proteins, are involved in these host-interaction processes upon infection [2]. PTMs comprise phosphorylation, acetylation, and methylation, as well as the incorporation of small polypeptides like ubiquitin or ubiquitin-related proteins such as the Small Ubiquitin-like Modifier (SUMO). It is known that a number of pathogens make advantage of PTMs for their own benefit; however, it is only established that a small number of pathogenic microorganisms may interfere with the SUMOylation pathways [2–6]. SUMOylation is a type of reversible post-translational modification that occurs in eukaryotic cells. In this regulation, a ubiquitin-like polypeptide called SUMO is strongly bound to the targeted proteins [7]. It is a crucial mechanism that regulates cellular functions such as the replication of DNA, the transcription of genetic information, the processing of RNA, and cell signaling [8,9]. Only lately have researchers begun looking into the strategies by which pathogenic bacteria alter SUMOylation in host proteins, and even now our understanding of these processes is limited [10]. In a recent study, we were able to show that the human pathogen *Staphylococcus aureus* inhibits the SUMOylation of host proteins in order to increase its intracellular survival and persistence for a longer period of time [11]. There is a correlation between the decreased degree of SUMOylation and the reduction in the amount of the SUMO-conjugating enzyme Ubc9. Moreover, over-expression of SUMO proteins in macrophages was shown to reduce bacterial intracellular proliferation, whereas treatment with ML-792, which blocked SUMOylation, led to an increase in bacterial load [11]. Interestingly, pathogens such as *Xanthomonas euvesicatoria* [12], *Yersinia pestis* [13], or *Listeria monocytogenes* [2] have been shown to release effectors that are able to elicit a general de-SUMOylation. *S. aureus* is a human pathogen that is extremely adaptable and is the cause of a variety of nosocomial and community-acquired infectious illnesses [14,15]. *S. aureus* is able to enter several different non-professional and professional phagocytes, where it may persist for many days [16–18]. The large number of virulence factors that *S. aureus* possesses is the primary reason for its prevalence as a pathogen and its capacity to be responsible for a broad range of disease manifestations [19,20]. PtpA is a low-molecular-weight protein tyrosine phosphatase that is secreted by *S. aureus*. We have previously shown that PtpA is released during growth and macrophage infection, and that deletion of *ptpA* reduces *S. aureus* intramacrophage survival and infectivity [21]. In this study, we show for the first time that a reduction in the levels of cellular SUMO-conjugated proteins is associated to the PtpA secreted virulence factor, which results in a reduction of Ubc9 protein level, the essential enzyme of the SUMOylation modification. In addition, we demonstrated that the phosphatase activity is required for the PtpA-dependent reduction in SUMOylation.

## 2. Materials and Methods

### 2.1. Bacterial strains and growth conditions

Strains and plasmids used in this study are listed in Table 1. Sequencing was used to confirm all mutant strains and plasmids used for this study. Strains of *Escherichia coli* were cultivated at 37°C in LB medium with the addition of 100 mg/ml ampicillin when needed. *S. aureus* isolates were either cultured in Tryptic Soy Broth (TSB; Becton Dickinson) medium at 37°C and 225 rpm with a culture to flask volume of 1:10 or plated on Tryptic Soy Agar (TSA; Becton Dickinson) medium supplemented with 10 mg/ml erythromycin when required. Bacterial growth in 96-well plates was observed using a microplate reader (Tecan, Lyon, France).

**Table 1.** Strains and plasmids used in this study

| Strain | Description <sup>1</sup> | Reference or source |
| --- | --- | --- |
| *S. aureus* |  |  |
| Newman | Laboratory strain (ATCC 25904), wildtype |  |
| Newman $\Delta ptpA$ | Newman $\Delta ptpA::lox66-aphAIII-lox71$; Kan<sup>R</sup> | [22] |
| Newman $\Delta ptpA::ptpA$ | Newman $\Delta ptpA$ derivative cis-complemented with pEC1_*ptpA*-Flag; Erm<sup>R</sup> | [22] |
| Newman $\Delta ptpA::ptpA\_D120A$ | Newman $\Delta ptpA$ derivative cis-complemented with pEC1_*ptpA\_D120A*-Flag; Erm<sup>R</sup> | This study |
| RN4220 $\Delta ptpA$ | RN4220 $\Delta ptpA::lox72$ | [22] |
| RN4220 $\Delta ptpA\_lox\_aph$ | RN4220 $\Delta ptpA::lox66-aphAIII-lox71$ | [22] |
| *E. coli* |  |  |
| IM08B | *E. coli* DC10B derivative harboring *hsdS* of *S. aureus* strain NRS384, $\Delta dcm$ | [23] |
| TOP10 | *E. coli* derivative ultra-competent cells used for general cloning | Invitrogen |
| **Plasmids** |  |  |
| pEC1 | pUC19 derivative containing the 1.45-kb *Clal erm(B)* fragment of Tn551 | [24] |
| pEC1_*ptpA*_D120A-Flag | pEC1 with a 1.4-kb fragment covering the *ptpA* ORF including a C-terminal flag tag and the aspartate 120 mutated to alanine, and 0.7-kb of the upstream region. | This study |

<sup>1</sup> Erm<sup>R</sup>, erythromycin-resistant; Kan<sup>R</sup>, kanamycin-resistant

### 2.2. Construction of the S. aureus ptpA cis-complementation strain Newman ptpA::ptpA_D120A

For *cis*-complementation of the *ptpA* mutation in strain Newman Δ*ptpA*, the vector pEC1_*ptpA* previously constructed [22] was used as template to generate the suicide plasmid *pEC1_ptpA_D120A* derivative by using the QuikChange Site-Directed Mutagenesis Kit (Agilent Technologies) with the primer #1559 (5’-GGAAGAGAGTGATGTACCAGCTCCATACTACACGAATAATT-3’). Plasmid *pEC1_ptpA_D120A* was then electroporated into the strain RN4220 Δ*ptp*, a marker-free *ptpA* variant of *S. aureus* strain RN4220, which was previously constructed [22]. The RN4220 derivative that integrated *pEC1_ptpA_D120A* was subsequently used as a donor for transducing the cis-integrated *pEC1_ptpA_D120A* into Newman Δ*ptpA*_lox_aph strain [22], thereby replacing the aph-tagged ptpA deletion with the *ptpA_D120A* derivative. Replacement of the *ptpA* deletion with the ptpA_D120A in Newman Δ*ptpA::ptpA* was confirmed by sequencing.

### 2.3. Macrophages culture and infection

The murine macrophage cell line RAW 264.7 (mouse leukemic monocyte macrophage, ATCC TIB-71), was cultured in Dulbecco’s modified Eagle’s medium (DMEM) (ThermoFisher Scientific), which was augmented with 10% foetal bovine serum and kept at 37°C in a humidified atmosphere containing 5% carbon dioxide. Lentiviral transduced Raw264.7 cell lines that expressed 6His-tagged SUMO1 and SUMO3 proteins were previously generated [11]. *S. aureus* was inoculated into the RAW 264.7 cells (5×10^5^ cells/mL, in 24 well plates) at a MOI of 20:1 (bacteria:cells), and the cells were then incubated for 1 h at 37°C and 5 % CO2. The residual extracellular bacteria were eliminated by incubating the cells with gentamicin (100 μg/mL) for 30 minutes after the cells had been washed once with PBS. Following gentamicin treatment, macrophages were twice washed with PBS (T0), and they were subsequently incubated 24 h in DMEM with 5 μg/mL lysostaphin. Macrophage lysates were diluted and plated on TSB agar plates and incubated at 37°C. The number of bacterial colonies at time post-gentamicin/number of bacterial colonies at T0 x 100 percent was used to calculate the survival rate of bacteria.

### 2.4. Immunoblotting

Infected macrophages were lysed in 100 μL of 2.5X Laemmli buffer, boiled for 10min at 95°C, sonicated for 10 sec at 50 % amplitude (DIGITAL Sonifier, Model 450-D, BRANSON) and centrifuged for 1min at 12000 x g. Proteins were separated on SDS-PAGEs, transferred to PVDF membranes, and analyzed by Western-blot using an anti-SUMO1 (#21C7, Developmental Studies Hybridoma Bank) or anti-SUMO2/3 antibody (#8A2, Developmental Studies Hybridoma Bank) as primary antibody and a HRP-coupled donkey-anti-mouse antibody as secondary antibody (Jackson ImmunoResearch, Interchim, France). The immunoblots were revealed with the Enhanced Chemiluminescence Detection kit (ChemiDocTM, Biorad) and quantified using Image Lab software (BioRad).

### 2.5. Quantitative RT-PCR (qRT-PCR)

Following the manufacturer’s instructions, total RNAs were extracted using the RNeasy^®^ plus Mini kit (Qiagen, GmbH, Germany). To measure the levels of mRNA expression, one microgram of total RNA was reverse-transcribed using the SuperScript III^®^ Reverse Transcriptase kit from Invitrogen. Using SYBR Green qPCR Master Mix (Roche) and specific primers (Table 2), quantitative RT-PCR (qRT-PCR) was carried out using a Light-Cycler 480 (Roche, France). As internal controls for mRNA quantification, mouse -actine was utilised. Using the Ct technique, the fold-induction was determined as follows: ΔΔCt = (Cttarget gene – Ctinternal control)treatment – (Cttarget gene – Ctinternal control)non-treatment, and the final data were derived from 2–ΔΔCt.

**Table 2.**
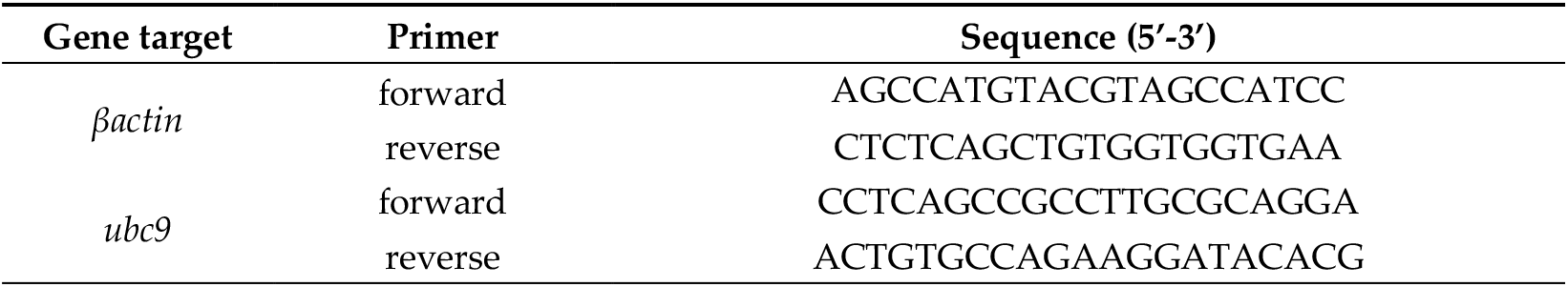
qRT-PCR primer used in this study

### 2.6. Statistical Analyses

The statistical significance of changes between groups was determined using the GraphPad software package Prism 9.4.0. *P* values < 0.05 were considered statistically significant.

## 3. Results

### 3.1. S. aureus PtpA phosphatase activity is required for intramacrophage survival

Since we have shown in the past that PtpA improves *S. aureus* surviving inside macrophages, we decided to assess wether the phosphatase activity of PtpA was necessary for the intramacrophage survival of *S. aureus* [21]. In order to test this, a *ptpA_D120A* mutant in *S. aureus* strain Newman was created using isogenic replacement. It was established previously that the residue D120 in the PtpA catalytic loop is necessary for the phosphatase activity of PtpA [22]. Next, we determined how the *S. aureus* strains Newman wild-type, Newman Δ*ptpA*, Newman Δ*ptpA*::*ptpA*, and Newman Δ*ptpA::ptpA_D120A* survived in RAW 264.7 cells 24 hours after being treated with gentamicin (pGt). The survival rate of the *ptpA* mutant had a substantial drop of around 50 percent when compared to the survival rate of macrophages that had been infected with the wild-type strain, while similar intracellular survival rates were observed in infected cells when the cis-complemented Newman Δ*ptpA::ptpA* strain was used (Fig. 1a). Therefore, these results are consistent with our prior intramacrophage survival findings confirming that PtpA plays an essential role in the infection process [21]. Macrophages that had been infected with the Newman Δ*ptpA::ptpA_D120A* mutant had, interestingly, a survival rate that was similar to the *ptpA* mutant (Fig. 1a). Based on these findings, we can conclude that the phosphatase activity of PtpA is required for the survival of *S. aureus* inside macrophages. In addition, we used the lactate dehydrogenase (LDH) assay to determine whether the lower survival of *S. aureus* mutant strains may be associated with a cytotoxic effect on the infected macrophage cells. The LDH test is a classic assay for identifying necrotic cell death by evaluating the level of damage to the cellular plasma membrane by measuring the amount of LDH enzyme that is released into the culture media. Infection with *S. aureus* strains did not elevate LDH discharge in culture supernatants over time in infected RAW 264.7 macrophages (Fig. 1b). This indicates that *S. aureus ptpA* derivatives are able to survive intracellularly for extended periods of time within macrophages without inducing cytotoxicity. Morevover, the possibility that the *ptpA_D120A* mutation would impair bacterial growth was ruled out by *in vitro* growth curves obtained with both wild-type and Newman derivative strains (Fig. 1c).

**Figure 1.**
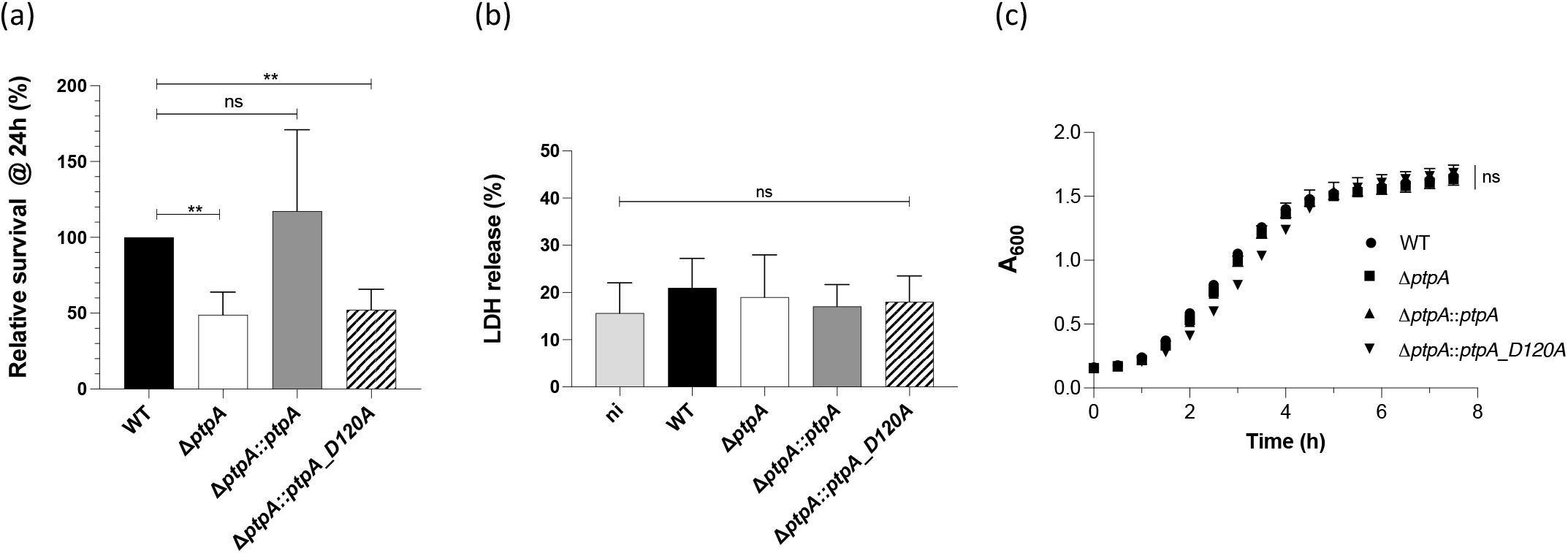
*S. aureus* PtpA derivative strains survival in macrophages. **(a)** Long-term survival of *S. aureus* in infected macrophages. *S. aureus* strains Newman wild-type (black bar), Newman *DptpA* (white bar), Newman Δ*ptpA::ptpA* (grey bar) and Newman Δ*ptpA::ptpA_D120A* (hatched bar) were infected, respectively, with RAW 264.7 macrophages at a MOI of 20 and coincubated for 1 h at 37°C before being treated with gentamicin/lysostaphin to get rid of non-phagocytosed bacteria. Bacteria were counted on plates after macrophages lysis with Triton X100 (0.1 %) at 24 hours post-pGt. The survival rates are expressed in relation to the number of intracellular bacterial cells counted just after Gentamicin administration. Data show means and standard deviations (n=4). **, *p*<0.01; ns, not significant (Mann-Whitney *U* test). **(b)** LDH release was evaluated using the CyQUANT test kit after macrophage cells were infected with bacteria at a MOI of 20 for 24 hours. The cells were seeded in a 96-well plate. The data are shown relative to the one hundred percent positive control on four biological replicates. ns, not significant (Mann-Whitney *U* test) **(c)** *In vitro* growth kinetics. Growth of *S. aureus* strains Newman wild-type (black symbols), Newman *DptpA* (white symbols), Newman Δ*ptpA::ptpA* (grey symbols) and Newman Δ*ptpA::ptpA_D120A* (hatched symbols) were performed in TSB at 37°C and 110 rpm in 96-well plates using a microplate reader (Tecan, Lyon, France). Data represent the mean A600 readings at the time points indicated (n=3). ns, not significant (One-way ANOVA test).

### 3.2. S. aureus PtpA phosphatase isn involved in the decrease of host SUMOylation upon infection

Given the fact that *ptpA* deletion or *D120* mutation reduces *S. aureus* intramacrophage survival, and that in our recent study we were able to show that *S. aureus* inhibits the SUMOylation of host proteins in order to increase its intracellular survival and persistence for a longer period of time in macrophages [11], we wondered if PtpA could be involved in host SUMOylation response to *S. aureus* infection. In order to test this hypothesis, we analyzed the amounts of SUMO1- and SUMO2/3-conjugated proteins in RAW 264.7 cells that have not been infected or are infected with *S. aureus* derivatives at long term infection (24h). First, in comparison to non-infected cells, macrophages that were infected with the *S. aureus* Newman wild-type strain showed a significant and specific decrease in the amount of SUMO1 (Fig. 2a) and SUMO2/3 (Fig. 2b) modified proteins. This was observed in accordance with what was previously described [11]. On the other hand, we found that the global pattern of SUMO-conjugated proteins was not decreased when infected with the Newman Δ*ptpA* or Δ*ptpA::ptpA_D120A* mutants, respectively (Fig. 2). However, decreased SUMOylation profiles were restored in infected cells when the cis-complemented Newman Δ*ptpA::ptpA* strain was used (Fig. 2). These data strongly suggest that PtpA plays a role in the decrease in SUMOylation observed with wild-type *S. aureus*.

**Figure 2.**
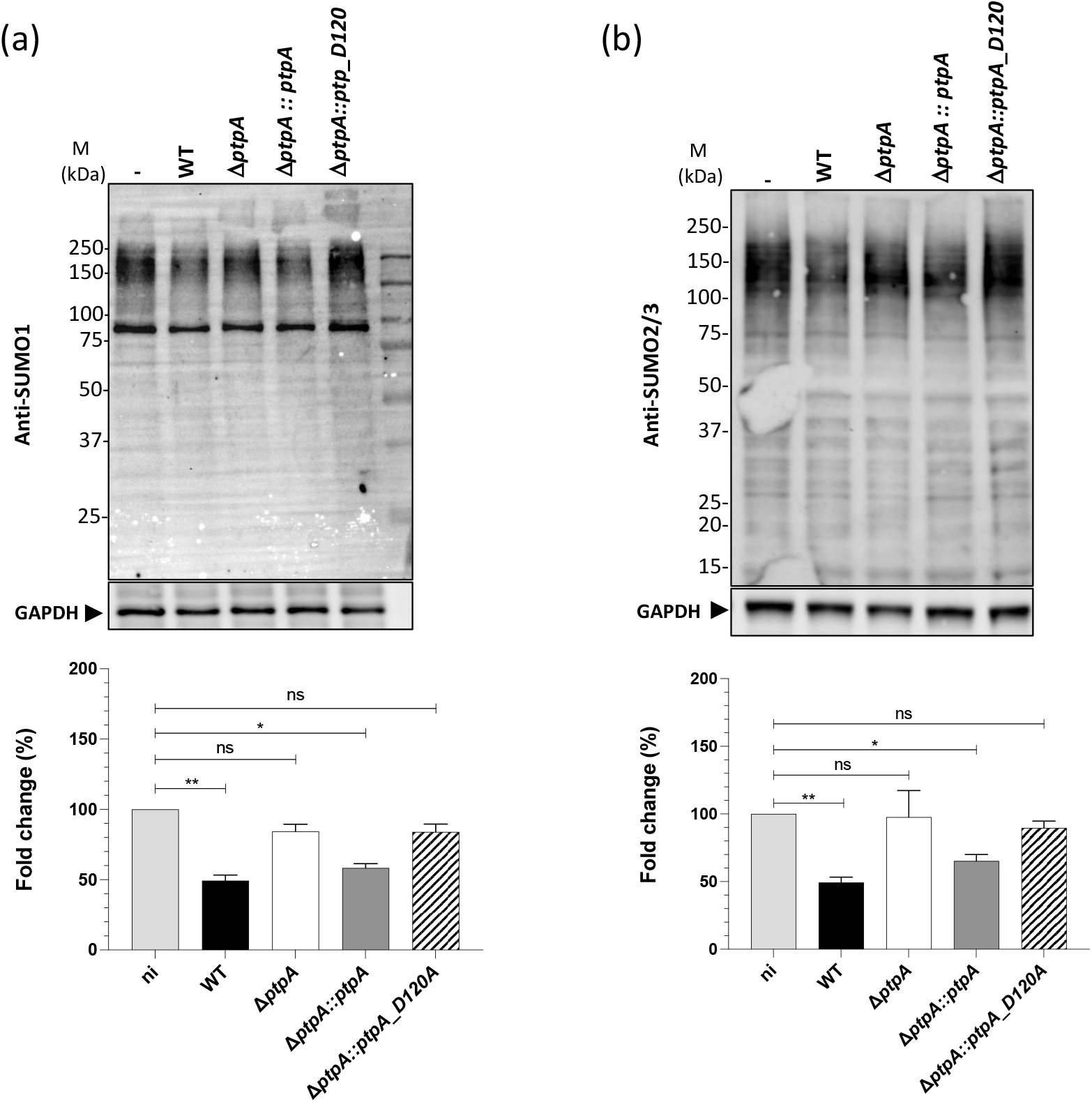
PtpA decreases SUMO-conjugated proteins in macrophages. Immunoblot analyses of the levels of SUMO1 **(a)**, SUMO2/3 **(b)**, and GAPDH in the lysates of macrophages infected with *S. aureus* strains for 24 h post-gentamycin treatment. Using Image Lab software (ChemiDoc), SUMO1 and SUMO2/3 smears were quantified from four different experiments and normalised to GAPDH (lower panels). The fold change chart displays the proportion of SUMOylated proteins recovered from infected cells in comparison to the quantity of SUMOylated proteins in non-infected control macrophages. The data represented are the mean ± SD of four independent biological experiments. **p < 0.01; *p < 0.05; ns, not significant (Kruskal-Wallis test followed by Dunn’s post hoc test).

### 3.3. S. aureus PtpA reduces Ubc9 level in a transcriptional-independent manner

After determining that PtpA is a critical virulence factor for *S. aureus* intracellular survival and that it is responsible for regulating the reduction of SUMOylation in macrophages, we wondered how PtpA might accomplish this regulation. The ubiquitin conjugating enzyme 9 (Ubc9), which is the only E2 conjugating enzyme in the SUMOylation machinery, is required for the SUMOylation procedure to occur [9]. Therefore, in order to determine how PtpA may interfere with the host’s SUMOylation machinery, we began by measuring the amount of the Ubc9 enzyme present in macrophages that had been infected with either wild-type or PtpA *S. aureus* derivative strains. Ubc9 protein level in infected macrophages with *S. aureus* wild-type or the *ptpA* complemented strain showed a decrease of approximately 50 percent at 24h of infection as compared to uninfected cells. However, no notable reduction was observed in macrophages infected with the Δ*ptpA* nor the inactive *ptpA_D120A* mutant strains (Fig. 3a). In addition, we utilised the proteasome inhibitor MG132 in order to evaluate whether or not the proteasome had a role in the reduction in the amount of Ubc9. The inhibition of proteasome activity by MG132 had no impact on the amount of Ubc9 (Fig. 3b), which continued to drop in infected macrophages 24 hours post-infection regardless of whether or not the *S. aureus* strains express PtpA. In addition, we used qRT-PCR to examine the levels of *ubc9* expression and found that *S. aureus* infection affects the expression of the Ubc9 enzyme at the mRNA level, although this variation occurs independently of PtpA. (Fig. 3c). These findings suggest that PtpA does, in fact, have an effect on the quantity of Ubc9, but that the reduction in this level is independent, respectively, of the proteasome and the transcriptional pathways.

**Figure 3.**
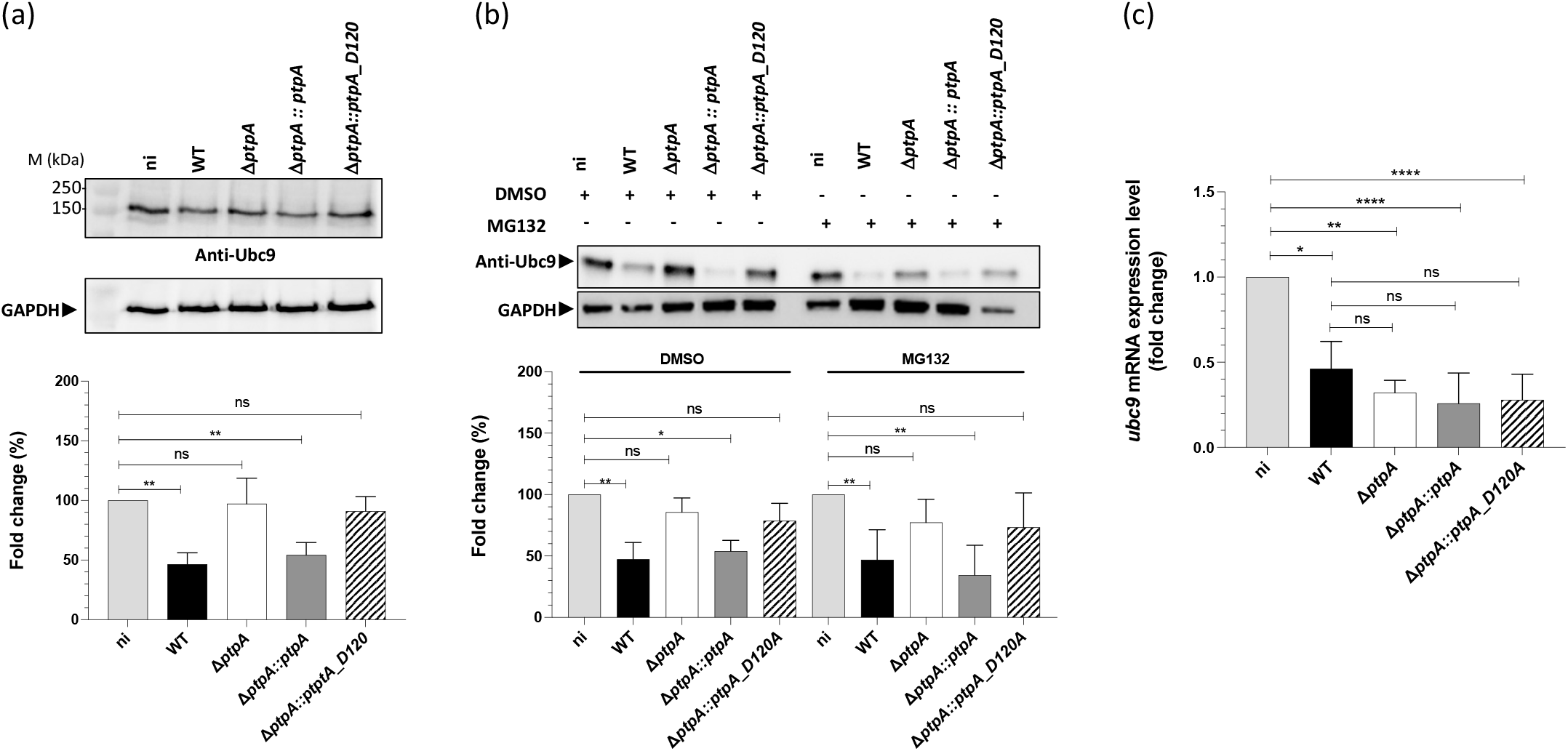
*S. aureus* PtpA reduces Ubc9 level but not *ubc9* transcription. Immunoblot analysis of Ubc9 and GAPDH levels in lysates of macrophages not treated **(a)**, or treated with MG132 for 3 hours prior to infection **(b)**, and infected with *S. aureus* strains for 24 h post-gentamicin treatment. n.i, noninfected control cells. Ubc9 bands were quantified from four independent experiments and normalized to GAPDH levels. The graph represents fold change compared to noninfected (lower panel). *p < 0.05; ns, not significant (Kruskal-Wallis test followed by Dunn’s post hoc test). **(c)** The influence of PtpA on the transcription of the *ubc9* gene. qRT-PCR was used to perform quantitative assessments of the *ubc9* transcript in *S. aureus* cells that had been cultured for 24 hours after being treated with gentamicin. Quantification of transcription rates was done in relation to the transcription of *βact* (in copies per copy of *βactin*), which was used as the standard. The data are provided as mean +SD of four separate biological experiments. *p < 0.05; ns, not significant (Mann-Whitney *U* test).

### 3.4. SUMOylation over-expression confirms the role of PtpA to promote intracellular survival of S. aureus

We used RAW 264.7 macrophages to artificially increase the level of SUMOylated proteins by over-expressing SUMO1 or SUMO3 enzymes in order to even further prove the role of PtpA in the host SUMOylation response to *S. aureus* infection. *S. aureus* derivatives were used to infect macrophages that over-expressed SUMO1 or SUMO3, and the quantity of viable intracellular bacteria was counted after 24 hours post-infection. Over-expression of either SUMO1 or SUMO2 decreased intracellular replication *S. aureus* wildtype and complemented *ptpA* strains compared to control macrophages expressing GFP (Fig. 4a), whereas Δ*ptpA* and *ptpA_D120A* mutant strains presented a considerably even lower survival rate. These results confirm, as previously observed, that an increase in SUMOylation in host cells has a negative impact on *S. aureus* ability to survive in the intramacrophage environment [11], but importantly demonstrate that PtpA expression is necessary to minimise the SUMOylation host response in order to improve *S. aureus* longterm survival. In addition, the function of PtpA in the regulation of *S. aureus* replication inside of macrophages that had been pretreated with an inhibitor of the SAE1/SAE2 enzyme was addressed [25]. Macrophages that were treated with the ML-792 inhibitor exhibited a substantial increase in the amount of *S. aureus* intracellular replication regardless of the strains used to infect the treated macrophages (Fig. 4b). According to these findings, treatment with the ML-792 inhibitor is able to circumvent the PtpA SUMOylation-dependent macrophage response.

**Figure 4.**
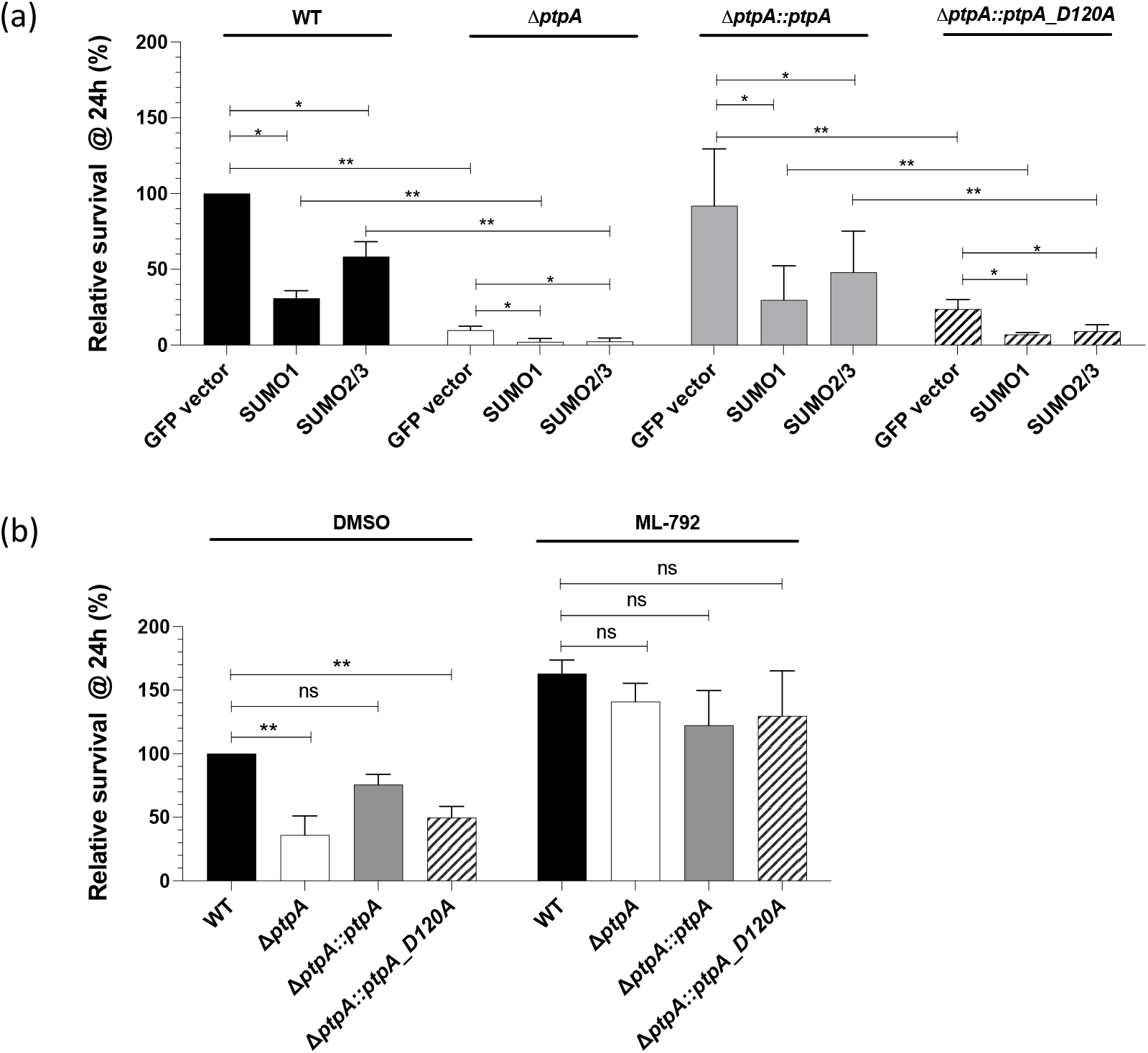
Impact of SUMOylation over-expression or inhibition on intracellular survival of *S. aureus ptpA* strain derivatives. **(a)** Intracellular *S. aureus* strains survival from macrophages overexpressing SUMO1 or SUMO2/3 *versus* control macrophages (GFP vector). Intracellular bacteria were counted after cell lysis and relative survival is presented as the ratio of intracellular bacteria at 24 h post-gentamicin compared to cells transfected with an empty GFP-vector, considered as 100%. (**b)** Macrophages pretreated with ML-792 at 0.5 μM or DMSO were infected with *S. aureus* strains. Number of intracellular bacteria recovered from macrophages 24 h post-gentamicin was counted and is presented as the ratio of intracellular bacteria compared to cells pretreated with DMSO, considered as 100%. **p < 0.01; *p < 0.05; ns, not significant (Mann-Whitney *U* test).

## 4. Discussion

Only lately have researchers looked into the processes that pathogenic bacteria use to modify the SUMOylation of host proteins, and even now our understanding of these mechanisms is limited [26,27]. Different pathogens have the ability to interfere with the host SUMOylation response by targeting different SUMO enzymes or pathways [27]. *Salmonella typhimurium*, an enteropathogenic bacterium, is responsible for a reduction in the SUMOylation host response. This is accomplished by the overexpression of two microRNAs that post-transcriptionally reduce Ubc9 expression [4]. *Shigella flexneri* is responsible for the modification of SUMO-conjugated proteins that are important in the regulation of mucosal inflammation and epithelial infiltration [3,28]. The adherent-invasive *Escherichia coli* (AIEC) bacterium might restrict autophagy by altering the host’s SUMOylation, thus enabling intracellular proliferation [5]. By reducing the induction of host inflammatory pathways, *Klebsiella pneumoniae* diminishes SUMOylation to enhance infection [6]. More recently, we have established that *S. aureus* promotes a reduction in macrophages SUMOylation response, thereby promoting its intracellular persistence [11].

Pathogens such as *Xanthomonas euvesicatoria* [12] and *Yersinia pestis* [13] have been shown to release effectors that are able to imitate host deSUMOylases, which in turn induces deSUMOylation of host proteins. Morevover, the pore-forming toxin listeriolysin (LLO) is responsible for *Listeria monocytogenes* ability to modify the host SUMOylation response by degrading Ubc9 [2]. However, the probable function of secreted effectors in the host SUMOylation response following *S. aureus* infection was unknown. In this study, we showed that (i) the secreted phosphatase PtpA was associated with the reduction of the macrophages SUMOylation response to enable *S. aureus* persistence at late stages of infection, (ii) that the PtpA phosphatase function was required in order to induce the SUMOylation response, (iii) that Ubc9 protein level was markedly decreased and PtpA-dependent, (iv) that the survival of the *S. aureus ptpA* mutant strains was significantly decreased by the over-expression of SUMO1 or SUMO3 in macrophage host cells, suggesting the involvement of PtpA in this SUMO-dependent regulation, and (v) that when macrophages were treated with the SUMOylation inhibitor, ML-792, *S. aureus ptpA* mutants were able to rescue PtpA deficiency and survive to a greater extend. According to our findings, PtpA is responsible for a global deSUMOylation in host cells at a later stage of infection by lowering the level of the Ubc9 protein to promote its intracellular survival. Nevertheless, treatment with the proteasome inhibitor MG132 had no effect on the amount of Ubc9 after *S. aureus* infection, demonstrating that Ubc9 degradation is independent of the proteasome. In addition, PtpA does not have any effect on the expression of Ubc9. Other virulence effectors that are released by bacterial pathogens have been demonstrated to downregulate Ubc9, which indicates that interfering with host SUMOylation via this critical enzyme of the SUMO machinery is a method that is shared by many different kinds of pathogenic bacteria, and now including *S. aureus* [27]. However, the PtpA-dependent target that is responsible for the reduction in Ubc9 protein level remains to be identified, and the specific mechanism of action to be defined.

As a result of the finding that a PtpA catalytically active tyrosine phosphatase is required to induce the host SUMOylation reduction, we came up with the hypothesis that a Tyr-phosphorylated-dependent control mechanism may be involved. Because the *ptpA* deletion mutant has no direct effect on the level of *ubc9* gene expression, post-translational regulation seems to be the most likely mode of regulation. One possible hypothesis would be that PtpA affects the phosphorylation status of Ubc9, which in turn influences the stability of the protein [29]. Due to the fact that PtpA is a tyrosine phosphatase and that Ubc9 is known to be phosphorylated on threonine residues [30,31], it is extremely unlikely that PtpA could directly dephosphorylate Ubc9. It is possible that the phosphorylation status of Ubc9 is influenced by a cascade of kinases that are phosphorylated on tyrosine residues. One putative candidate could be the Ser/Thr protein kinase Akt that is activated through tyrosine phosphorylation [32]. Moreover, the Akt protein directly phosphorylates Ubc9 at Thr35 leading to the formation of Ubc9 thioester bond [30]. If this is the case, one hypothesis would be that PtpA, could dephosphorylate these regulatory kinase, which in turn would have an indirect effect on the level of Ubc9. However, such PtpA Tyr-phosphorylated substrates remain to be identified to date.

In conclusion, the current work showed for the first time that the secreted phosphatase PtpA is capable of reducing the Ubc9 conjugation enzyme level to impede host SUMOylation response, thus promoting *S. aureus* survival at long-term infection. SUMOylation crosstalk during bacterial infection represents a promising area of research that will not only further our knowledge of how SUMOylation occurs in cells, but may also reveal potential targets for therapeutic treatment against *S. aureus* infections and persistence at long-term infections.

## Author Contributions

Conceptualization, V.M. and L.G.-Z.; methodology, V.M. and L.G.-Z.; formal analysis, V.M., L.G.-Z., and N.Y.; investigation, N.Y., M.M., P.S. and M.B.; resources, V.M. and L.G.-Z; writing—original draft preparation, V.M., M.B., P.S. and L.G-Z. All authors have read and agreed to the published version of the manuscript..

## Informed Consent Statement

Not applicable.

## Data Availability Statement

The datasets generated during and/or analyzed during the current study are available from the corresponding author on reasonable request.

## Acknowledgments

The authors thank Noémie Quelin et Nabila Sebbagh for excellent technical support.

